# Cortical state and natural movie responses in cat visual cortex

**DOI:** 10.1101/031765

**Authors:** Martin A. Spacek, Nicholas V. Swindale

**Author notes:** Conflict of Interest: The authors declare no competing financial interests.

## Abstract

**Abstract:** How does cortical state affect neural responses to naturalistic stimuli, and is it analogous between anesthetized and awake animals? We recorded spikes and local field potential (LFP) from all layers of isoflurane-anesthetized cat primary visual cortex (V1) while repeatedly presenting wide-field natural scene movie clips. Spiking responses of single units were remarkably precise, reliable and sparse, with lognormally distributed mean firing rates. Many units had distinct barcode-like firing patterns, with features as little as 10 ms wide. LFP-derived cortical state switched spontaneously between synchronized (1/*f*) and desynchronized (broadband). Surprisingly, responses were more precise, reliable and sparse during the synchronized than desynchronized state. Because the desynchronized state under anesthesia is thought to correspond to attending periods in awake animals, during which responses are enhanced, our results complicate the analogy between cortical states in anesthetized and awake animals. The presence of orientation maps in cat V1 may explain contrary reports in anesthetized rodents, and predicts a similar result in anesthetized ferret and primate V1.

**Significance Statement:** Global brain activity changes spontaneously over time and can be characterized along a spectrum from slow synchronized activity, to fast desynchronized activity. This spectrum is similar in awake, asleep and anesthetized animals, but is its effect on neural responses the same in all cases? Here we show that neural responses to natural movies in anesthetized cat visual cortex are more precise during synchronized activity. This is contrary to reports in anesthetized rodents, which we suggest may be due to greater columnar organization in cat visual cortex. Since this is also contrary to enhanced responses and behavioural performance during attention, when activity is desynchronized, our results suggest that similar brain states in awake and anesthetized animals may not be functionally analogous.

## Introduction

As a complex dynamic system, the brain is never in exactly the same state twice. Spontaneous changes in brain state were noted in even the earliest electroencephalograms(Berger, 1929). However, most studies that repeatedly present identical stimuli implicitly assume that the brain is in the same state at the onset of each trial, and that averaging over trials will provide a reasonable estimate of response variability. This may not always be the case, even under anesthesia (Arieli et al., 1996; Petersen et al., 2003). Brain state can influence response variability,and should therefore be accounted for.

Brain state can be characterized along a spectrum from synchronized to desynchronized (Destexhe et al., 1999; Harris and Thiele, 2011). The synchronized state consists of large amplitude low frequency fluctuations, and occurs during deep anesthesia,slow-wave sleep, and awake quiescent periods. The synchronized state can be further subdivided into up and DOWN phases (Destexhe et al., 1999; Sanchez-Vives and McCormick, 2000; Harris and Thiele, 2011), corresponding to periods of higher and lower resting membrane potential. The desynchronized state consists of low amplitude high frequency fluctuations, and occurs during light anesthesia, rapid eye movement sleep, and awake attending behavior.

Visual neuroscience has traditionally relied on reduced stimuli such as drifting bars and gratings to characterize neural responses. Naturalistic stimuli can elicit responses that are poorly predicted from responses to reduced stimuli (Olshausen and Field, 2005). Although reduced stimuli have much lower dimensionality than naturalistic stimuli, relying too heavily on reduced stimuli may obscure insights into how the brain processes visual information. To more fully characterize neural populations in visual cortex, it is therefore important to also consider responses to naturalistic stimuli. Natural image sequences are spatially naturalistic, but the gold standard is natural scene movies, which are both spatially and temporally naturalistic.

How precise and reliable are natural scene movie responses in V1], andhow does cortical state influence them? We examined single unit responses across all layers ofV1 in isoflurane-anesthetized cats, while stimulating with natural scene movies containing saccade-like camera movements. Cortical state varied spontaneously over time, and was characterized by the frequency content of deep-layer LFP. Recordings were divided into synchronized and desynchronized periods. Spiking responses to natural scene movies were remarkably precise, reliable and sparse,consisting of barcode-like patterns of response events consistent across trials, some as little as 10 ms wide. Signal and noise correlations were weak overall (∼ 0.1 and 0.02) at the 20 ms time scale, but were stronger in the synchronized than desynchronized state. Contrary to reports in primary sensory cortices of anesthetized rodent (Goard and Dan, 2009; Marguet and Harris, 2011; Hirata and Castro-Alamancos, 2011; Zagha et al., 2013; Pachitariu et al., 2015), natural scene movie responses in anesthetized cat V1 were more precise, reliable and sparse in the synchronized than desynchronized state. In the synchronized state, trial-averaged responses were also better correlated with motion within the movie. This is surprising, because the synchronized state under anesthesia is analogous to quiescent periods in awake animals and the desynchronized state to alert attending periods, and neural responses are known to be enhanced to attended stimuli (Roelfsema et al., 1998; Fries et al., 2001; Cohen and Maunsell, 2009; Mitchell et al., 2009; Chalk et al., 2010).

Our results therefore complicate the analogy between cortical states in anesthetized and awake animals. A possible explanation for conflicting reports in primary sensory cortices of anesthetized rodents may be that cat V1 has orientation maps,which rodent V1 lacks. Standing and travelling waves (Petersen et al., 2003; Massimini et al., 2004; Benucci et al., 2007; Luczak et al., 2007; Xu et al., 2007; Mohajerani et al., 2010; Sato et al., 2012) of activation (up phases) in the synchronized state may therefore interact differently with incoming stimuli in V1 of higher mammals. This explanation predicts a similar result in anesthetized V1 of other species with orientation maps, such as ferrets and primates.

## Materials and Methods

### Surgical procedures

Animal experiments followed the guidelines of the Canadian Council for Animal Care and the Animal Care Committee of the University of British Columbia. After initial sedation, cats were intubated and mechanically ventilated (Harvard Apparatus, Holliston, MA) at ∼ 20 breaths/min to maintain end-tidal CO_2_ of 30-40 mmHg. Anesthesia was maintained by inhalation of 0.5-1.5% isoflurane with 70% N_2_O in O_2_. Blink and pinna (ear) reflexes and toe pinch were used to ensure sufficient anesthetic depth. During surgical procedures and euthanization, up to 3% isoflurane was used. Intramuscular injection of dexamethasone (1 mg/kg) was used to reduce swelling and salivation. The animal was hydrated by intravascular (**IV**) infusion of a mixture of lactated Ringer’s salt solution(10-20 mL/h), sometimes with added potassium chloride (20 mEq/L) and dextrose (2.5%). Heart rate and blood oxygenation were monitored with a pulse-oximeter (Nonin 8600V), with the sensor placed on the tongue or a shaved portion of tail. Mean arterial blood pressure was monitored with a doppler blood pressure monitor(Parks Medical 811-B) on a shaved section of hind leg. Body temperature was maintained at 37°C via closed-loop control with a homeothermic blanket (Harvard Apparatus). All vital signs were logged during the course of each experiment. Experiments lasted up to 3 days each.

Animals were placed in a stereotaxic frame on an air table, with ear bars coated in topical anesthetic (5% lidocaine). Local anesthetic (bupivacaine) was injected subcutaneously around the top of the skull and into the ear muscles before cutting the skin to expose the skull. A roughly 4 × 6 mm craniotomy (1-5 mm lateral and 3-9 mm posterior relative to the centerline and earbar zero, respectively) was drilled with a dental drill (Midwest Stylus, DENTSPLY Professional, Des Plaines, IL) over Brodmann’s area 17 and 18. A stereo surgical microscope was used during drilling, removal of meninges, and polytrode insertion. Artificial cerebrospinal fluid (ACSF) was used to flush away blood and other detritus from the meninges, and to keep them moist. Ophthalmic surgical sponges (Ultracell Eye Spears, Aspen Surgical, Caledonia, MI) were used to wick blood and excess fluid away. Care was taken to not apply pressure to the brain. A small area of dura was dissected away one layer at a time with an ophthalmic slit knife (Beaver Optimum 15°, BD Medical, Le Pont-de-Claix, France; or ClearCut 3.2mm, Alcon, Mississauga, ON). A small nick in the pia was then made with the ophthalmic slit knife to allow for polytrode insertion. Prior to insertion, CSF was wicked away from the point of insertion using an ophthalmic surgical sponge to improve unit isolation. Immediately before or after insertion, high purity low temperature agarose (Type III-A, Sigma-Aldrich, St. Louis, MO) dissolved in ACSF at a concentration of 2.5-4% was applied in liquid form at 38-40°C to the craniotomy. This quickly set and eliminated brain movement due to heart beat and respiration. The polytrode was advanced perpendicular to the pial surface using a manual micromanipulator (Model 1460 Electrode Manipulator, David Kopf Instruments, Tujunga, CA) under visual control, until the topmost electrode sites disappeared below the surface of the cortex. Any further advancement was made with a hydraulic micromanipulator (Narishige MHW-4, East Meadow, NY), typically 150- 300 **μ**m at a time.

Nictitating membranes were retracted with phenylephrine (10%, 1-2 drops/eye), and pupils were dilated with tropicamide (0.5%, 1-2 drops/eye).Custom-made rigid gas permeable contact lenses (14 mm diameter, 7.8-8.7 mm base curvature, +2.00 to +4.00 diopter, Harbour City Contact Lens Service, Nanaimo, BC) protected the eyes and refracted the cat’s vision to the distance ofthe stimulus display monitor. To improve focus, 3 mm diameter artificial pupils were placed directly in front of the lenses. To prevent eye drift, one animal (ptc22, Fig. 1 & Fig. 2d,e) was given an initial IV bolus ofthe systemic paralytic pancuronium bromide (1 mg/kg), and paralysis was maintained by constant rate infusion (0.2 mg/kg/h). For the other two animals (ptc17 & ptc18, Fig. 2a–c) *α*- bungarotoxin was instead injected retrobulbarly (125 *μ*M, 0.5 mL per eye) as a local paralytic. Eye position was closely monitored by reverse ophthalmoscopy to ensure stability, using fine blood vessels as landmarks. Receptive fields (mapped online with a manually controlled light or dark bar) fell within a few degrees ofthe area centralis.

### Recordings

Extracellular recordings were made from cortical area 17 of 3 anesthetized adult cats (2 male, 1 female), using 54-site single shank (15 *μ*m thick, 207 *μ*m wide, 1138 or 1325 *μ*m long) silicon polytrodes (Blanche et al., 2005) (NeuroNexus, Ann Arbor, MI), with electrode sites arranged in 2 or 3 columns in a hexagonal layout (50 or 65 *μ*m spacing). Results presented here include data from 5 recording sessions, 3 in the left hemisphere and 2 in the right. These 5 recording sessions were from 4 unique hemispheres in 3 cats, for a total of 47.8 hours of recording. Histological track reconstruction was not successful.

Extracellular voltage waveforms from all 54 electrode sites were unity-gain buffered by a pair of 27-channel headstages (HS-27, Neuralynx, Tucson, AZ), and amplified by a 64-channel 5000 × amplifier with fixed analog filters (FA-I-64, Multichannel Systems, Reutlingen, Germany). The first 54 channels of the amplifier were high-pass analog filtered (0.5-6 kHz) for use as spike channels. Data from a subset of 10 of the 54 electrode sites, evenly distributed along the length of the polytrode, were also separately low-pass analog filtered (0.1-150 Hz) for use as LEP channels. All 64 channels were then digitally sampled (25 kHz for the high-pass channels, 1 kHz for the low-pass channels) by a pair of12-bit 32-channel acquisition boards with an internal gain of 1-8× (DT3010, Data Translations, Marlboro, MA), controlled by custom software written in Delphi (Blanche et al., 2005).

### Spike sorting

Spike sorting was done using custom open source software written in Python (http://spyke.github.io). A “divide-and-conquer” spike sorting method (Swindale and Spacek, 2014) translated correlated multisite voltages into action potentials of spatially localized, isolated neurons. This method tracked neurons over periods of many hours despite drift, and distinguished neurons with mean firing rates < 0.05 Hz. Briefly,the steps in this method were: 1) Nyquist interpolation to 50 kHz and sample-and-hold delay correction (Blanche and Swindale, 2006); 2) spike detection; 3) initial clustering based on the channel of maximum amplitude; 4) spike alignment within each cluster; 5) channel and time range selection around the spikes in each cluster; 6) dimension reduction (multichannel PCA, ICA, and/or spike time) into a 3D cluster space; 7) clustering in 3D using a gradient-ascent based clustering algorithm (GAC) (Swindale and Spacek, 2014); 8) exhaustive pairwise comparisons of each cluster to every other proximal cluster, generally involving multiple iterations of steps 4-7. Each spike was localized in 2D physical space along the polytrode by fitting a 2D spatial Gaussian to the signal amplitudes using the Levenberg-Marquardt algorithm. Free parameters were *x* and *y* coordinates, and spatial standard deviation. To improve detection of low firing rate units, spike sorting was performed on entire recording sessions lasting up to 12 hours each, even though only a small subset of each session was relevant to this study.

### Visual stimulation

Visual stimuli were presented with millisecond precision using custom open source software written in Python (http://dimstim.github.io) based on the VisionEgg library (Straw, 2008)(http://visionegg.org). Stimuli were displayed on a flat 19” (36 × 27 cm) CRT monitor (Iiyama **HM903DTB**) at 800 × 600 resolution and 200 Hz refresh rate. A high refresh rate was used to prevent artifactual phase locking of neurons in V1 to the screen raster (Williams et al., 2004). One recording(ptc17.tr2b.r58, Fig. 2a) intentionally used a low 66 Hz refresh rate in an attempt to induce phase-locking, but this did not affect the results presented here. The monitor was placed 57 cm in front of the cat’s eyes. At this distance, 1 cm on the screen subtended 1° of visual angle, and the monitor subtended horizontal and vertical angles of ˜ 36° and 27° respectively. The monitor had a maximum luminance of 116 cd/m^2^. Display monitors are typically gamma corrected to linearize output light levels when presenting computer-generated stimuli such as bars and gratings. However, gamma correction was not applied here during natural scene movie presentation because gamma correction already occurs in cameras during the video capture process (Poynton, 1998).

Movies were acquired using a hand-held consumer-grade digital camera (Canon PowerShot SD200) at a resolution of 320×240 pixels and 60 frames/s. Movies were filmed close to the ground, in a variety of wooded or grassy locations in Vancouver, BC. Footage consisted mostly of dense grass and foliage with a wide variety of oriented edges. Focus was kept within 2 m and exposure settings were set to automatic. The horizontal angle subtended by the camera lens (51.6°) was measured for proper scaling to match the visual angle subtended by the movie on the stimulus monitor. Two example movie clips are available at http://dimstim.github.io: `MVI̲ 1400̲ 200-500`, corresponding to Fig. 2a, c; and MVI̲ 1403̲ 0- 300 corresponding to Fig. 1 and the upper panels of Fig. 3 & Fig. 4. Others are available upon request. Movies contained simulated saccades (peaks in Fig. 11a) of up to 275^°^/s, generated by manual camera movements in order to mimic gaze shifts (eye and head movements), which can exceed 300°/ s in cat (Munoz et al., 1991). The movies contained little or no forward/backward optic flow. Movies were converted from color to grayscale, and were presented at 66 frames/s. Depending on the refresh rate (see above), each frame corresponded to either 1 or 3 screen refreshes. Global motion was calculated for every neighboring pair of movie frames (Farnebäck, 2003) using the OpenCV library (http://opencv.org). Global contrast and luminance were calculated for each frame by taking the standard deviation and mean, respectively, of all the pixel values in each frame. Other stimulus types were also presented during each recording session, including drifting bars and gratings, flashed gratings, m-sequence white noise movies, and blank screen, but these were not included in this study.

### Cortical state characterization

Cortical state was determined from the deep-layer LEP. First, 60 Hz mains interference was digitally filtered out using a 0.5 Hz wide elliptic notch filter (negative peak in Fig. 2f). The spectrogram was then constructed by dividing the signal into 2 s wide overlapping time bins at 0.5 s resolution, applying a Hanning window, and calculating the fast Fourier transform separately for each time bin. The synchrony index (SI) was defined as the L/(L+H) ratio (Saleem et al., 2010), calculated as a function of time from the deep-layer LEP spectrogram using 30 s wide overlapping time bins at 5 s resolution. L and H are the power in low (0.5-7 Hz) and high(15-100 Hz) LFP frequency bands, respectively. SI ranged from 0 to 1, with 1 representing maximum synchronization. SI thresholds for the synchronized and desynchronized state were > 0.85 and < 0.8, respectively. However, visual inspection of the spectrogram was used in tandem with the SI, so the above thresholds were not hard limits. Choosing a lower SI threshold for the desynchronized state to limit analysis to desynchronized periods with a more consistent LEP spectrum did not substantially change results (not shown).

### Response characterization

Spike and LEP analyses were performed using custom open source software (Spacek et al., 2009) written in Python (http://neuropy.github.io). Each unit’s peristimulus time histogram (PSTH, i.e., the response averaged over trials) was calculated by convolving a Gaussian of width 2*σ* = 20 ms with the spike train collapsed across all trials that fell within the recording period of interest. This timescale was chosen because 20 ms is roughly the membrane time constant of neocortical layer 5 (Mainen and Sejnowski, 1995) and hippocampal **CA1** (Spruston and Johnston, 1992) pyramidal neurons. This is also the timescale at which hippocampal pyramidal cell spike times are best predicted by the activity of peer neurons, and therefore may bethe most relevant for cell assemblies (Harris et al., 2003). The analyses shown in Fig.6 were repeated for a range of 2*σ* values (10-100 ms), and the conclusions were independent of the precise value chosen.

Detecting response events in a trial raster plot is a clustering problem: how do spike times cluster together into response events, with temporal density significantly greater than background firing levels? As for spike sorting (see above), spike time clustering was performed using the GAC algorithm (Swindale and Spacek, 2014), witha characteristic neighborhood size of 20 ms. Spike time clusters containing less than 5 spikes were discarded. The center of each detected cluster of spike times was matched to the nearest peak in the PSTH. A threshold of *θ* = *b* + 3 Hz was applied to the matching**PSTH** peak, where *b* = 2 median(*x*) is the baseline of each **PSTH** *x*. Peaks in the PSTH that fell below *θ* were discarded, and all others were treated as valid response events. The equation for *θ* was derived by trial and error, and visual inspection of all 1870 detected peaks in all 563 **PSTH**s confirmed that there were no obvious false positive or false negative detections. This threshold for detecting peaks in the **PSTH**s did not cause a sudden cutoff at the low end in thenumber of spikesper detected response event per trial (Fig. 8a). The precision of a response event was defined as its width, measured as the temporal separation of the middle 68% (16th to 84th percentile) of spike times within each cluster.

Response reliability was quantified as the mean pairwise correlation of all trial pairs of a unit’s single trial responses (Goard and Dan, 2009). Single trial responses were calculated by dividing single trial spike trains into 20 ms wide overlapping time bins at 0.1 ms resolution, and counting the number of spikes within each time bin. This resulted in a matrix of integer values as a function of time, with one row per trial. Pearson’s correlation was calculated between all possible pairs of trials. For trial pairs in which one or both trials had no spikes, their correlation was set to 0. The reliability of each cell during each cortical state was defined as the mean of all of the pairwise correlations of the trials during that state. Response reliability could range from —1 to 1, but was mostly positive.

The sparseness S of a signal, whether **PSTH**, absolute value of LEP, or MUA, was calculated

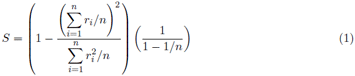

where *r_i_* ≥ 0 is the signal value in the *i^th^* time bin, and *n* is the number of time bins (Vinje and Gallant, 2000). Sparseness ranges from 0 to 1, with 0 corresponding to a uniform signal, and 1 corresponding to a signal with all of its energy in a single time bin.

Although the 1 s period of blank screen separating each trial is shown at the end of each recording trace in Fig. 3–Fig. 5 & Fig. 10a, d, precision, reliability and sparseness measures in Fig. 6 & Fig. 10 excluded this inter-trial period of blank screen. The mean firing rate of each unit in a given cortical state (Fig. 8b) was calculated by its spike count in that state, divided by the state’s duration. Mean firing rates therefore included the 1 s period of blank gray screen between movie clip presentations. Units were not required to surpass a mean firing rate threshold for inclusion for analysis. The only requirement was that a unit was responsive, i.e., that it had at least one detected response event in its **PSTH**.

Multiunit activity (MUA) (Fig. 10d–f) was calculated by combining the spike trains of all isolated single units and convolving the resulting multiunit spike count signal with a Gaussian of width 2*σ* = 20 ms. MUA coupling was calculated by correlating each unit’s **PSTH** with the trial-averaged MUA excluding that unit. MUA coupling was calculated somewhat differently from the original method (Okun et al., 2015) by taking Pearson’s correlation between each **PSTH** and the MUA.

## Results

### Cortical state

Cortical state was characterized by the frequency content of the deep-layer LEP (Fig. 1). The synchronized state had large amplitude low frequency fluctuations with an approximately 1/ *f* distribution, while the desynchronized state consisted of lower amplitude fluctuations spanning a wider range of frequencies (Fig. 1a, b). Spontaneous transitions between the two states were visible in the LEP spectrogram (Fig. 1c). A synchrony index (SI) (Fig. 1d, **Materials and Methods**) was used to quantify the degree of synchronization over time. SI ranged from 0 to 1, where 1 represents maximum synchronization. The distribution of SI from all recordings is shown in Fig. 1d (inset). Based on both visual inspection of the LEPspectrogram and application of thresholds to the corresponding SI, recordings were divided into periods of synchronized, desynchronized and undefined states. Six natural scene movie recordings (3.5 h total duration, 5 penetrations in 3 cats) exhibited an obvious spontaneous change in cortical state (5 from desynchronized to synchronized, 1 from synchronized to desynchronized, Fig. 1c & Fig. 2a—e). A similar amount of time was spent in both states (104 min synchronized, 93 min desynchronized, 10 min undefined). A total of 219 single units were isolated in these 6 recordings.

### Natural scene movie responses

Spike raster plots of 3 example single units are shown in Fig. 3, in response to 400 presentations of two different wide-field natural scene movie clips, each 4.5 s in duration. One spontaneous cortical state transition occurred during each movie. Spike raster plots across trials exhibited a pattern reminiscent of **UPC** barcodes, consisting of remarkably precise, reliable and sparse response events. For both natural scene movies, this pattern was visibly more pronounced during the synchronized than desynchronized state. Each unit’s **PSTH** was classified as responsive during a given cortical state if it contained at least one response event. Response events were detected using an automated method to cluster spike times (**Materials and Methods**). Example **PSTH**s are shown underneath the raster plots in Fig. 3 & Fig. 5, with colored dots marking detected response events. A total of 267 out of a possible 563 **PSTH**s were classified as responsive. There were more responsive **PSTH**s in the synchronized than desynchronized state (153 vs. 114, χ^2^ test, *p* < 0.02), and significantly more response events in the synchronized than desynchronized state (1167 vs. 703, χ^2^ test, *p* < 7.4 × 10^−27^).

The 3 example units in Fig. 3 were responsive to both natural scene movie clips, but some units in that pair of recordings were responsive to only one movie and not the other. Fig. 4 shows 3 such example units. For the two natural scene movie recordings shown in Fig. 3 & Fig. 4, 51% (20/ 39) of responsive units were responsive during only one movie: 8 responded only to the first movie, and 12 responded only to the second. However, 50% (39/ 78) of units isolated in that penetration did not respond to either movie. Some units were responsive in one cortical state but nonresponsive in the other (Fig. 4b, g, Fig. 5c). Across all 6 recordings, 30% (49/ 163) of responsive units were responsive only during the synchronized state, 6% (10/ 163) were responsive only during the desynchronized state, and 64%(104/ 163) were responsive during both states.

**Figure 1.**
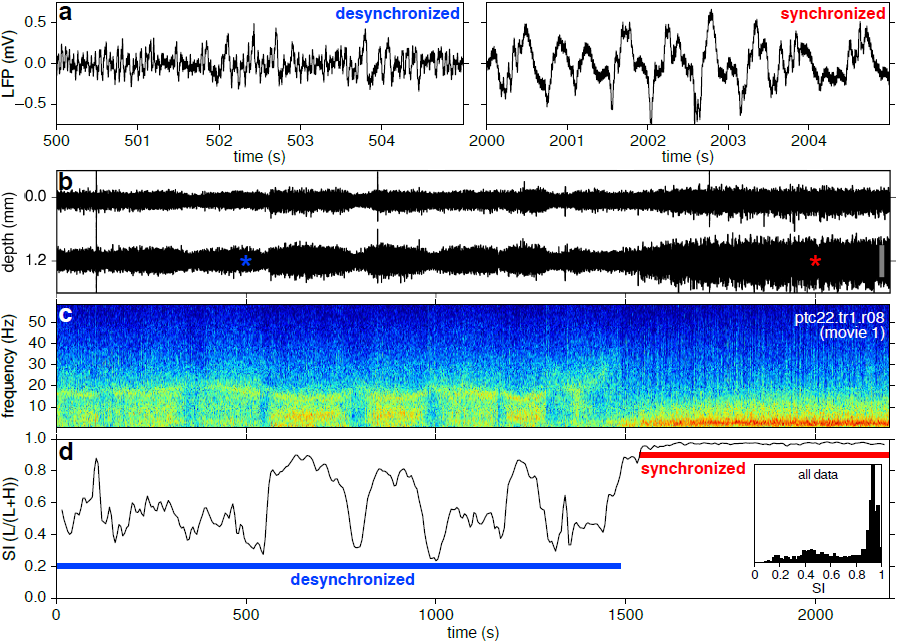
A spontaneous change in cortical state during 37 min of repeated presentation of a 4.5s natural scene movie clip. (**a**) Short representative deep-layer LFP voltage traces during the desynchronized and synchronized state. (**b**) Full duration superficial and deep-layer LFP, with depth measured from the top of the polytrode. Colored asterisks indicate time periods of the panels in (**a**). Scale bar: 1 mV. (**c**) Deep-layer LFP spectrogram. Red represents high power, blue low power (arbitrary units). The synchronized state had a ˜ 1/ *f* frequency distribution, while the frequency distribution of the desynchronized state was more broadband and variable.(**d**) Synchrony index (SI) calculated from the L/(L+ H) frequency band ratio of the spectrogram. Cortical state switched spontaneously from desynchronized to synchronized about 2/ 3 of the way through the recording. Blue and red horizontal lines indicate the duration of the desynchronized and synchronized periods, respectively. Inset, SI histogram for all 3.5 h of natural scenemovie recordings.

**Fig. 2.**
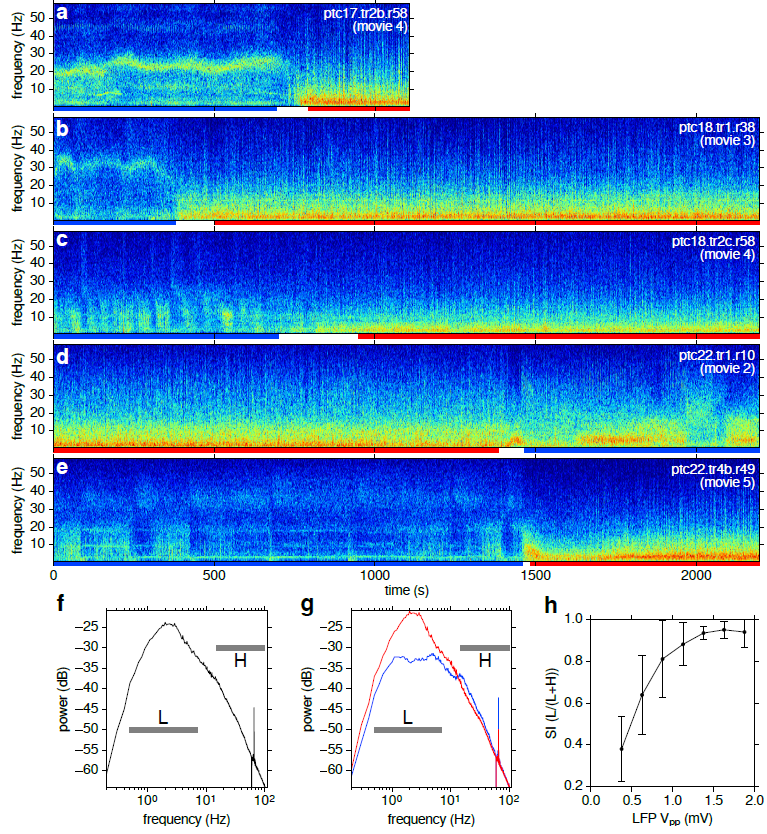
(**Previous page**) LFP spectrograms and power spectral density (PSD). (*a-e*) Spectrograms from 5 of the 6 recordings (in addition to that shown in Fig. 1c) during 200 (**a**) or 400 (**b-e**) presentations of a 4.5 s natural scene movie clip. Blue and red horizontal lines underneath each spectrogram indicate the duration of the desynchronized and synchronized periods, respectively, in each ofthe recordings, as determined from the SI (not shown). (**f**) PSD of all 6 recordings. Power is in decibels relative to 1 mV^2^. Horizontal lines mark the limits ofthe low (L) and high (H) bands used to calculate SI. On this log-logscale, thelow band is roughly centered on the broad peak at ˜ 2 Hz. Some of the attenuation below 1 Hz is due to analog filtering during acquisition. The narrow positive peak at 66 Hz corresponds to the movie frame rate, and the narrow negative peak at 60Hz is from filtering out mains interference (**Materials and Methods**).(**g**) Same as (**f**) but split into synchronized (**red**) and desynchronized (**blue**) periods,showing greater low frequency power in the synchronized state. (**h**) SI (mean ±1 standard deviation) covaried positively with LFP peak-to-peak amplitude (V_pp_, 0.25 mV wide bins).

The responses of another 3 example units to a different movie in a different cat are shown in Fig. 5. Even though the spectrogram and the SI in the desynchronized state was more consistent in this recording (Fig. 2b; Fig. 5a) than in the other two example recordings (Fig. 1c & Fig. 2d; Fig. 3a & e), responses for these example units were again visibly more precise, reliable and sparse in the synchronized than desynchronized state.

Response amplitude, precision, reliability and sparseness are summarized in Fig. 6 for all 267 units with at least one response event, across all 6 recordings during which a spontaneous change in cortical state occurred. All four measures were significantly greater in the synchronized than desynchronized state (means, *p* values, and statistical tests reported in Fig. 6). Five unique movie clips were presented in these 6 recordings. Response event amplitude was quantified as the height (in Hz) above baseline of each peak in the **PSTH** (**Materials and Methods**). Response event width (in ms) was quantified as twice the standard deviation of the spiketimes belonging to the event. Response reliability was quantified as the mean pairwise correlation of all trial pairs of a unit’s responses. The sparseness (Eq. 1) of each **PSTH** ranged from 0 to 1, with 0 corresponding to a uniform signal, and 1 corresponding to a signal with all of its energy in a single time bin.

There was no strong dependence of response precision, reliability and sparseness on unit position along the length of the polytrode (Fig. 7). Because polytrode insertions were generally vertical, and were inserted to a depth relative to the surface of the cortex (**Materials and Methods**), position along the polytrode roughly corresponded to cortical depth. In both cortical states, response precision and sparseness (Fig. 7a, c), but not reliability (Fig. 7b), were greater in superficial layers.

**Fig. 3.**
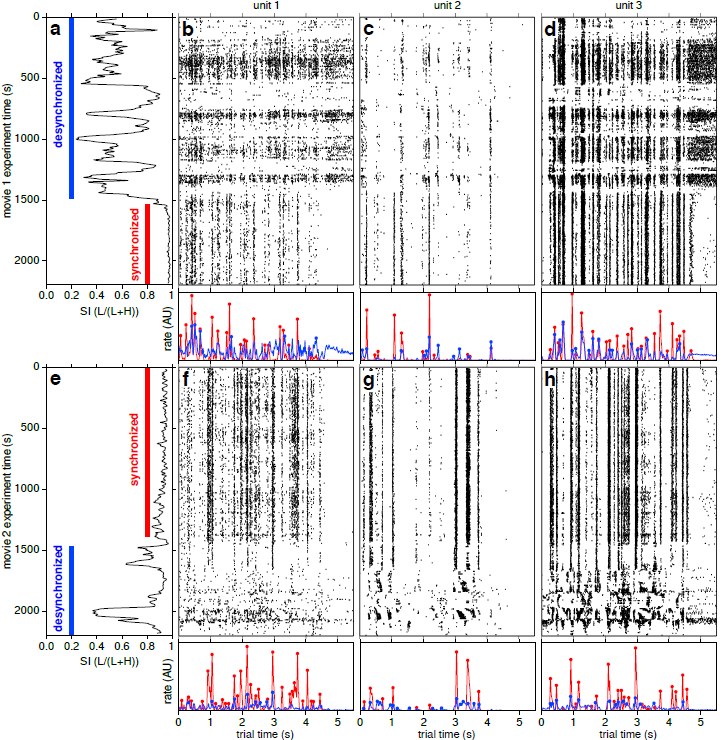
(**Previous page**) Cortical state affects precision, reliability and sparseness of natural scene movie responses. During 400 presentations (vertical axis) of two different 4.5 s (horizontal axis) natural scene movie clips (upper and lower panels) inthe same penetration, two spontaneous cortical state transitions occurred: from desynchronized to synchronized (**a**, same recording as in Fig. 1), and from synchronized back to desynchronized (e, same recording as in Fig. 2d). SI is shown in the leftmost column. Vertical colored lines indicate the duration of each cortical state (**red**: synchronized; **blue**: desynchronized). (**b-d**, **f-h**) Trial raster plots of natural scene movie responses of 3 example units (one per panel column), left to right in order of increasing depth from the top of the polytrode (161, 186 and 820 *μ*m, respectively). Each black tick represents one spike. Each presentation was separated by 1 s of blank gray screen (from 4.5 to 5.5 s of trial time). PSTHs are shown underneath each raster plot, color-coded by state, with dots marking detected response events. For display purposes, each PSTH panel uses a different vertical scale. For all 3 example units during both movies,responses were visibly more precise, reliable and sparse during the synchronized state than the desynchronized state. A 20 minute gap of blank gray screen stimulation separated the end of the first recording (**a**) from the start of the second (**e**). Patterns of response events were distinct for all 3 example units, even for the first two whose physical separation was only ˜ 25 *μ*m. AU: arbitrary units.

### Bursting and mean rates

Are the response events described above due to bursting, in which a single unit fires multiple spikes in close succession, or are they usually composed of no more than asingle spike on any given trial? The distributions of spike counts per response event per trial are shown in Fig. 8a, separately for each state. In both states, the distribution was very close to lognormal (dashed curves), with geometric means of 0.5 spikes/ event/ trial, well below 1 spike/ event/ trial. In the synchronized and desynchronized states, 78% and 76%, respectively, of response events had ≤ 1 spike/ trial. Therefore, > 75% of response events in either state were unlikely to be the result of bursting.

How might mean firing rates vary as a function of cortical state? Although intuition suggests that rates should be higher in the desynchronized state, previous reports show no clear relationship between mean firing rates and cortical state (Goard and Dan, 2009; Harris and Thiele, 2011). The mean firing rate of each unit during a cortical state was calculated by taking its spike count during that state and dividing by the duration of the state. The distributions of mean firing rates across the population are shown separately for both states in Fig. 8b. Mean firing rates spanned a wide range (0.0005-50 Hz), with a distribution that was approximately lognormal (dashed curves). This was the case in both states. Mean rates in the synchronized and desynchronized state were not significantly different(Mann-Whitney U test, ensemble geometric means of 0.18 and 0.14 Hz and standard deviations of 1.0 and 1.1 orders of magnitude, respectively).

### Correlations and MUA coupling

By definition, pairwise correlations of averaged single unit responses (signal correlations) and of trial-to-trial variability (noise correlations) should be greater in the synchronized than desynchronized state (Harris and Thiele, 2011). Signal correlations were calculated by taking Pearson’s correlation between **PSTH**s of all simultaneously recorded pairs of responsive single units. This was done separately for both cortical states. Similarly, noise correlations were calculated by taking Pearsonߣs correlation of the differencebetween eachunit’s single trial response and **PSTH**, for all pairs of single units, for both cortical states. Signal correlations were weakly positive on average, and were indeed significantly greater in the synchronized than desynchronizedstate (0.18 and 0.11, respectively, Mann-Whitney U test, Fig. 8c). Noise correlations were even weaker, but still positive on average, and significantly greater in the synchronized state (0.031 vs. 0.015, Fig. 8d). Signal and noise correlations in both stateshad a weak but significantly negative dependence on unit pair separation (Fig. 8e, f).

**Fig. 4.**
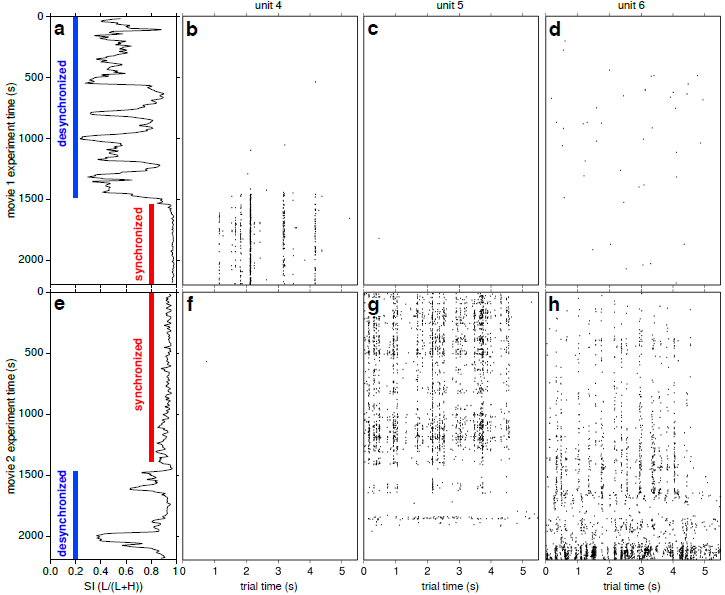
Same as **Fig. 3** (excluding **PSTH**s) but with 3 more example units, each of which had response events during one movie but not the other. Panels (**c**) & (**f**) had only one spike each. Two of the example units(**b,g**) had response events only during the synchronized state. Left to right, units are in order of increasing depth from the top ofthe polytrode (77, 974 and 1197 *μ*m, respectively). Although difficult to see in this layout, visual inspection revealed that the last two units in the second recording (**g,h**) shared several response events that fell within a few ms of eachother.

**Figure 5.**
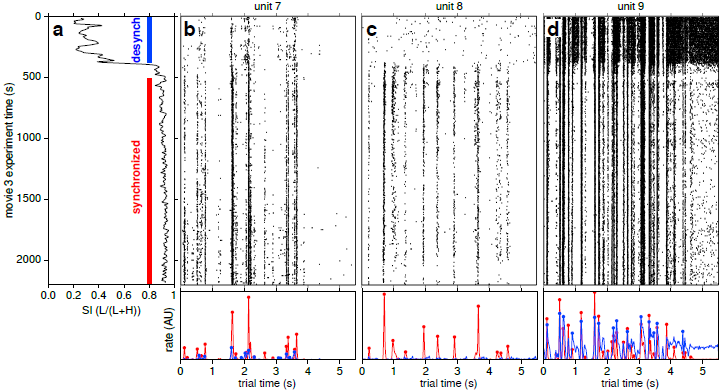
Responses of 3 more example units in a different recording in a different cat, to 400 presentations of a different movie clip (same layout as upper panels in Fig. 3). (**a**) SI over the course of 37 min of repeated presentation of a 4.5 s natural scene movie clip (same recording as in Fig. 2b). SI in the desynchronized state was more consistently low in this recording than in Fig. 3 & Fig. 4, yet the results were similar: responses were again visibly more precise, reliable, and sparse in the synchronized than desynchronized state. Left to right, units are in order of increasing depth from the top of the polytrode (367, 847 and 974 *μ*m, respectively). Again, although difficult to see in this layout, visual inspection revealed that the first and last units (**b,d**) shared several response events that fell within a few ms of each other, despite high physical separation (˜ 610 *μ*m). Neither unit shared any response events with the middle unit (**c**).

**Figure 6.**
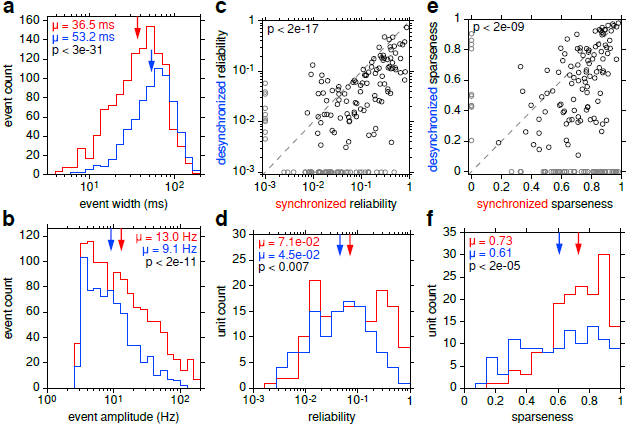
Response precision, reliability and sparseness vs. cortical state for all 6 recordings. (**a**)Distributions of response event widths during the synchronized (**red**) and desynchronized (**blue**)state. (**b**) Distributions of event amplitudes relative to baseline firing. (**c**) Scatter plot of response reliability in the two cortical states for all units that were responsive in at least one state. For display purposes, units with noresponse events during a cortical state were assigned a reliabilityof 10_−3_ in that state (**gray**). Significantly more units fell below the dashed *y* = *x* line than above it (83%, 136/ 163, p < 2 ⨯ 10_−17_, χ _2_ test). (**d**) Responsereliability distributions for thepoints in(c), excluding those set to 10_− 3_. (**e**) Scatter plot of response sparseness inthe two cortical statesfor all units that were responsive in at least one state. For display purposes, unitswith no responseevents during a cortical state were assigned a sparseness of 0 in that state. Significantly more units fell below the dashed *y* = *x* line than above it (74%, 120/163, p < 2 × 10_−9_, χ _2_2 test). (**f**)Response sparseness distributions for the points in (**e**), excluding those set to 0. Arrows denote geometric means in (**a**), (**b**) & (**d**), and arithmetic means in (bold). Response events were significantly narrower and higher, and responses were significantly more reliable and sparse in the synchronized than desynchronized state (p values in (**a**), (**b**), (**d**) & (**f**), Mann-Whitney U test).

**Figure 7.**
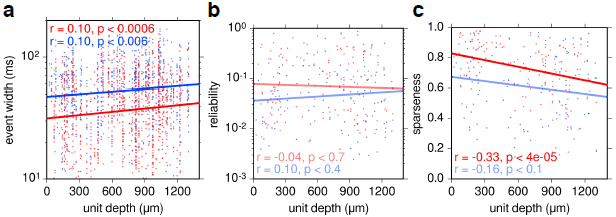
Single unit response precision, reliability and sparseness vs. unit depth from the top of the polytrode, for all 267 responsive units in all 6 recordings. (**a**) Each point represents a response event. Response event width was weakly but significantly positively correlated with unit depth in both the synchronized (red) and desynchronized (**blue**) state. Response precision was therefore weakly but negatively correlated with unit depth in both states. The difference in mean event width between the two states was consistent (˜ 16 ms) as a function of unit depth. (**b**) Each point represents a responsive **PSTH**. Response reliability was not significantly correlated with unit depth in either state. (**c**)Response sparseness was significantly negatively correlated with unit depth in only the synchronized state. Lines show least squares linear regression (two-sided Student’s T-test, *r*- and *p*- values shown in each panel). Desaturated lines and statistics denote insignificant correlations.

A recent report has shown that the degree of coupling between single unit and multi-unit activity (MUA) is a simple but consistent metric for characterizing single units, and that it can be used to predict both single unit signal correlations and the degree of synaptic connectivity with other neighboring neurons (Okun et al., 2015). How might MUA coupling relate to cortical states and natural scene movie responses in cat V1? MUA coupling was calculated for each single unit by calculating the trial-averaged MUA (e.g., Fig. 10d) from all single units, excluding the singleunit of interest, and correlating that with the unit’s **PSTH** (**Materials and Methods**). Thiswasdone for all single units during both cortical states. Fig. 9a shows the distributions of MUA coupling across the population. MUA coupling was significantly greater in the synchronized than desynchronized state (Mann-Whitney U test, *p* < 6 × 10^− 5^). Single unit response reliability was significantly and positively correlated with MUA coupling, in bothcortical states (Fig. 9b). However, response sparseness was not significantly correlated with MUA coupling in either state (Fig. 9c).

**Figure 8.**
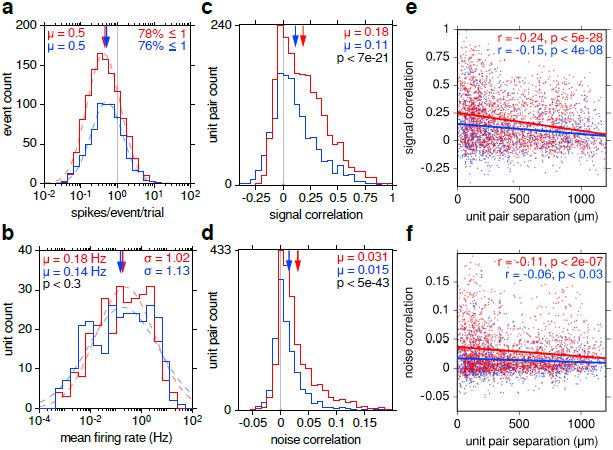
Response event spike counts, single unit mean firing rates, and correlations as a function of cortical state. (**a**) Distributions of the number of spikes per response event, per trial, for both cortical states (**red**: synchronized, **blue**: desynchronized). In both states, > 75% of response events averaged less than 1 spike per trial (vertical grey line), and were therefore not involved in bursting. Lognormal functions were fit to both distributions (dashed curves,Levenberg-Marquardt algorithm). Arrows denote geometric means (*μ*). (**b**) Mean firing rate distributions of all isolated single units. Distributions in the synchronized (285 **PSTH**s) and desynchronized (278 **PSTH**s) state were not significantly different from each other (Mann-Whitney U test, *p* < 0.3). Arrows denote geometric means. Standard deviations (**σ**) are expressed in powers of 10. Lognormal functions were fit to both distributions(dashed curves). (**c,d**) Distributions of signal and noise correlations for all responsive unit pairs in both states. Arrows indicate means. Correlations were on average weakly positive in both states, but significantly higher in the synchronized state (Mann-Whitney U test). (**e,f**) Signal and noise correlations vs. unit pair separation. Both types of correlations decreased slightly but significantly with increasing unit separation (mostly in depth) in both cortical states. Lines show least squares linear regression (two-sided Student’s T-test, *r*- and *p*-values shown).

**Figure 9.**
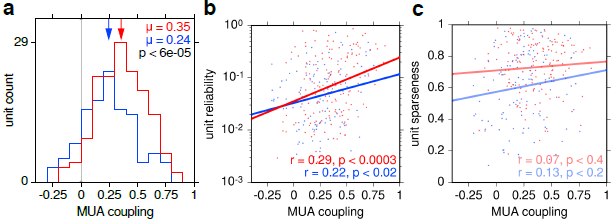
MUA coupling as a function of cortical state. (**a**) MUA coupling (the correlation of each single unit **PSTH** with the MUA, excluding that unit) distributions for all responsive units in the synchronized (**red**) and desynchronized (**blue**) states. MUA coupling was significantly greater in the synchronized state (Mann-Whitney U test, *p* < 6 × 10^− 5^). (**b,c**) Single unit response reliability and sparseness vs. MUA coupling for all responsive units. Single unit response reliability was significantly and positively correlated with MUA coupling, in both states, but sparseness was not. Lines show least squares linear regression (two-sided Student’s T-test, *r*- and *p*-values shown in each panel). Desaturated lines and statistics denote insignificant correlations.

### LEP and MUA reliability and sparseness

Given that single unit responses during natural scene movie stimulation were more reliable and sparse in the synchronized state (Fig. 6), does the same hold for the LFP and MUA? Trial-aligned LEP and MUA are shown in Fig. 10a, d in both cortical states for one example recording. As expected, the amplitudes of the LFP and MUA were greater in the synchronized state (shown more explicitly for LFP in Fig. 2h). LFP and MUA reliability were measured in a similar way as for singleunit responses, using Pearson’s correlation between the signal on each trial and the mean of the signal on all other trials. This was done for all trials in both states in all 6 recordings (988 desynchronized trials, 1093 synchronized trials). LFP and MUA reliability were both significantly greater in the synchronized than desynchronized state (Fig. 10b, e). The sparseness of each LEP and MUA trace was also measured (for LFP, sparseness of the absolute value of the signal was used). Response sparseness was also significantly greater in the synchronized state (Fig. 10c, f).

### Stimulus representation

How do precise and reliable single unit responses, such as those shown in Fig. 3- Fig. 5, relate to the visual stimulus, and how does stimulus representation vary with cortical state? Calculating receptive fields from short repetitive natural scene movie clips is a difficult and perhaps intractable problem, given the spatial and temporal correlations inherent to movies (Carandini et al., 2005), and the low number of movie frames per clip (300 for each of the 5 unique clips used here). Instead, responses were compared to the global motion, contrast and luminance calculated as a function of time from all of the on-screen pixels of each movie clip (**Materials and Methods**). The correlation between each responsive unit’s **PSTH**and movie global motion, contrast and luminance signals was calculated separately in each cortical state. Fig. 11a shows movie frames and the global motion signal of an example movie clip (`MVI̲ 1400̲ 200-500`, **Materials and Methods**), as well as the **PSTH** of an example single unit in both cortical states. Movie clips consisted of simulated saccades generated by manually rotating the camera with short, quick motions. This resulted in a highly kurtotic distribution of global motion within the movies (Fig. 11b). The correlation between responsive **PSTH**s and global motion was weakly positive, and significantly greater in the synchronized than desynchronized state (Fig. 11c, d, mean values of 0.091 and 0.041 respectively). This was when calculated at a delay of 30 ms (2 movie frames) between stimulus and response. The mean **PSTH**-motion correlation as a function of stimulus-response delay is shown in Fig. 11e. Not only was it greatest in the synchronized state at adelay of 30 ms, but stimulus-response delay modulated **PSTH**-motion correlation more in the synchronized than desynchronized state. In comparison, single unit responses were much more weakly correlated with global movie contrast and luminance (taken as the standard deviation and mean, respectively, of the pixel values of each frame), and did not differ significantly as a function of cortical state (Fig. 12). However, both contrast and luminance were again more strongly modulated as a function of stimulus-response delay in the synchronizedthan desynchronized state (Fig. 12c, f).

**Figure 10.**
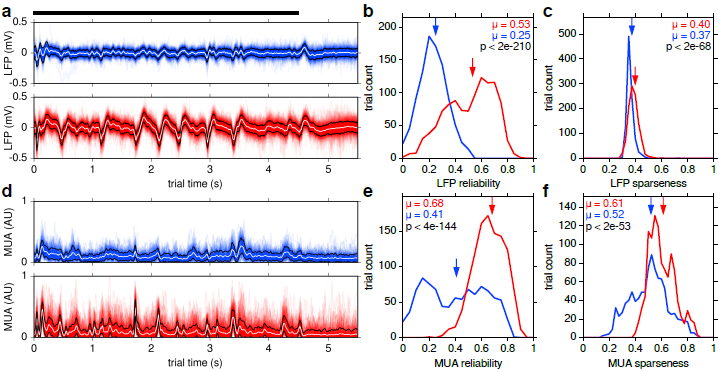
Trial-aligned LEP and multi-unit activity (MUA) were more reliable and sparse in the synchronized state. (**a**) Trial-aligned deep-layerLEP traces are shown as semi-transparent lines, in the desynchronized (**blue**, 127 trials) and synchronized (**red**, 227 trials) state, for an example recording (same as Fig. 2c). Mean ±1 standard deviation are shown as white and black lines, respectively. Black horizontal bar represents movie clip duration. (**b**) Distributions of LFP trial reliability (Pearson’s correlation between the LFP of each trial and the mean of the LFP of all other trials),for both states in all recordings. (**c**) Distributions of the sparseness ofthe absolute value of the LFP of each trial, for both states in all recordings. (**d— f**) Same as (**a— c**) but for MUA, calculated by combining spike trains from all isolated single units (**Materials and Methods**). All distributions were significantly higher in the synchronized than desynchronized state (Mann-Whitney U test, *p* values shown in each panel). Arrows indicate means. Bin widths are 0.05 in (**b**) & (**e**) and 0.025 in (**c**) & (**f**).

The sudden global motion of a movie saccade is highly salient, and may be enough to simultaneously depolarize many cells and induce an up phase, during which spike timing may be more precise (Luczak et al., 2007). Since there are typically multiple movie saccades per trial (Fig. 11a), this might reset the state of the neural population at multiple time points within each trial. In thesynchronized state, up and DOWN phases are better separated in time (Luczak et al., 2013), anda movie saccade might therefore more reliably trigger an up phase in the synchronized than desynchronized state. The presence of movie saccades might therefore be a somewhat trivial explanation for greater response precision in the synchronized than desynchronized state (Fig. 6). This hypothesis predicts that as the elapsed time since the last movie saccade increases,the precision of response events should decrease. Although response events were indeed more likely to occur shortly after a movie saccade than at other times (Fig. 13a), the above prediction did not hold: response precision was only very weakly correlated with time since the last preceding movie saccade, and significantly so only in the desynchronized state (Fig. 13b). In addition, response sparseness was insignificantly correlated with movie motion sparseness (Fig. 13c). Movie saccades are therefore not likely responsible for the greater precision and sparseness of responses in the synchronized than desynchronized state.

**Fiure 11.**
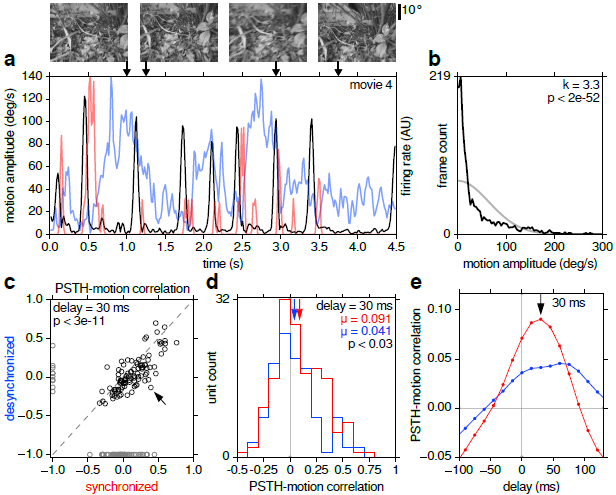
Global motion within movies and its effect on responses. (**a**) Movie frames(**top**) and global motion amplitude (**bottom, black**) for one example movie. Motion peaks correspond to sudden camera movements, approximating saccades and head movements. The **PSTH** of one example unit is shown in the synchronized (**red**) and desynchronized (**blue**) state. Allowing for stimulus-response delay, **PSTH** peaks for this example unit tracked motion amplitude better in the synchronized state. (**b**) Distribution of motion amplitude for all 5 unique movie clips (**black**). Bin widths are 4 deg/s wide. The distribution was highly kurtotic (*k* = 3.3), significantly moreso than a normal distribution (Anscombe-Glynn kurtosis test, *p* < 2 × 10^− 52^). A normal distribution with the same standard deviation and probability mass is shown for comparison (gray). (**c**) Scatter plot of correlation between global motion and responsive **PSTH**s 30 ms later, in the desynchronized vs. synchronized state. For display purposes, units that were nonresponsive in a given state were assigned a value of —1 (**gray**). Excluding these, significantly more units fell below the dashed *y* = *x* line than above it (83%, 86/ 104, *p* < 3 × 10^−11^, χ^2^ test). Arrow denotes the example unit shown in (**a**). (**d**) Distribution of the points in (**c**) in the synchronized (**red**) and desynchronized (**blue**) state, excluding points assigned a value of —1. Arrows denote means. **PSTH**-motion correlations were significantly higher in the synchronized state (Mann-Whitney U test, *p* < 0.03). (**e**) Mean **PSTH**-motion correlations in both states as a function of delay between stimulus and response. **PSTH**-motion correlations peaked at 30 ms and were more strongly modulated by delay in the synchronized state.

**Figure 12.**
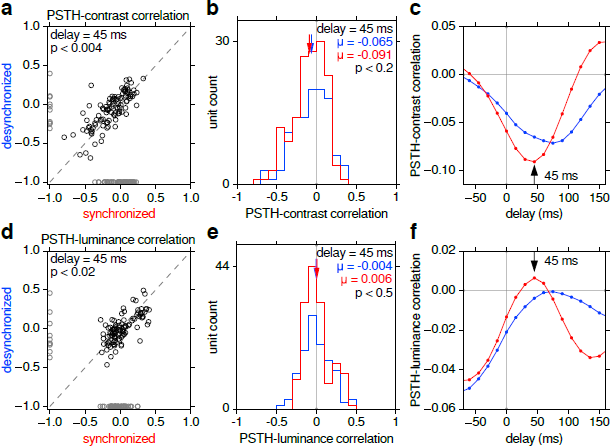
Same as Fig. 11c–e, but for movie contrast (**top**) and luminance (**bottom**)instead of motion. Unlike motion, neither showed significantly different correlations with single unit responses as a function of cortical state. (**a,d**) Scatterplots of correlation between responsive **PSTH**s and global contrast and luminance for desynchronized vs. synchronized states. (**a**) At 45 ms delay, fewer units (**black**) fell below the dashed *y* = *x* line than above it (36%, 37/ 104, *p* < 0.004, χ^2^ test). (**d**) At 45 ms delay, more units fell below the dashed *y* = *x* line than above it (62%, 64/104, *p* < 0.02, χ ^2^ test).For a significance threshold of *p* = 10^−6^, neither χ^2^ test was significant, while that in Fig. 11c was. (**b,e**) Distributions corresponding to (**a,d**). In both cases, means were not significantly different between the synchronized (**red**) and desynchronized (**blue**) state (Mann-Whitney U test, *p* values shown). (**c,f**) Mean **PSTH**-contrast and **PSTH**-luminance correlations in both states as a function of stimulus-response delay, which peaked at 45 ms and 60 ms, respectively. Both were more strongly modulated by delay in the synchronized state, as was the case for **PSTH**-motion correlations (Fig. 11e).

**Figure 13.**
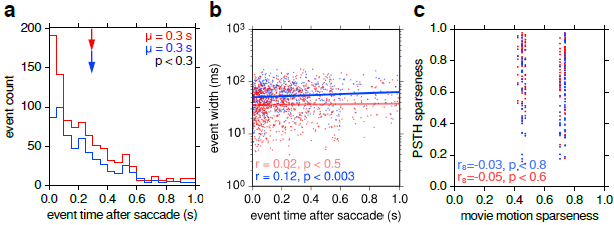
Response precision is not dependent on time since the nearest preceding movie saccade. (**a**) Response events were more likely to occur immediately following a movie saccade, in both the synchronized (**red**) and desynchronized (**blue**) state. The average time of a response event following the nearest preceding movie saccade (0.3 s) was not significantly different between the two states (*p* < 0.3, Mann-Whitney U test). (**b**) Response event width was only very weakly correlated with time since the preceding movie saccade, and significantly so only in the desynchronized state. This is the opposite of what would be expected if responses were more reliable during the synchronized state due to more effective resetting of the network from movie saccades. Lines show least squares linear regression (two-sided Student’s T-test) (**c**) Response sparseness was not significantly correlated with movie motion sparseness in either state (Spearman’s rank-order correlation, two-sided Student’s T-test).

## Discussion

Single unit responses to natural scene movie clips consisted of barcode-like response events (Fig. 3–Fig. 5), some as little as 10 ms in duration (Fig. 6a). Across the population of units, there was great diversity in the patterns of response events, as shown by the low mean pairwise signal correlations between units (Fig. 8c). There was also a surprisingly wide range of mean firing rates, most below 1 Hz, which approximately followed a lognormal distribution (Fig. 8b), inlinewith an increasing number of reports in various species and cortical areas (Wohrer et al., 2013; Buzsáki and Mizuseki, 2014). Interestingly, the distribution of spike counts per response event per trial was also lognormal (Fig. 8a), and low enough to preclude bursting as a major component of response events.

There are a handful of reports of temporally precise, reliable and sparse responses to natural scene movies in V1: in awake behaving macaque (Vinje and Gallant, 2000), and in anesthetized cat, both extracellularly (Yen et al., 2007; Herikstad et al., 2011) and intracellularly (Haider et al., 2010; Baudot et al., 2013). Similar precision and reliability have been reported in awake behaving macaque area MT during random dot stimulation with low motion coherence (Bair and Koch, 1996). There have been more reports of even greater temporal precision (events as little as ∼ 1 ms wide) of responses to high-entropy stimuli in retinal ganglion cells (RGCs) of salamander and rabbit (Berry et al., 1997), and in the lateral geniculate nucleus (LGN) of anesthetized cat (Alonso et al., 1996; Reinagel and Reid, 2000).

As visual information propagates from RGCs to LGN to V1, the temporal precision and reliability of responses generally decrease (Kara et al., 2000). It is interesting to consider that this precision is retained at all. LGN inputs constitute only a small fraction of synapses onto (mostly layer 4) cortical cells, yet these inputs are very effective at driving the cortex (Ahmed et al., 1994). Convergent event-like input from LGN during naturalistic stimulation may be another reason for this strong drive (Alonso et al., 1996; Wang et al., 2010). There may be an evolutionary benefit in maintaining temporal precision in V1. Sparse coding (Olshausen and Field,1996) and the energy efficiency (Attwell and Laughlin, 2001) that comes with it may beone such reason. Another may relate to delay line coding, which proposes that precise relatively-timed spikes allow for simple scale-invariant representations of stimuli (Hopfield, 1995). This theory is supported by increasing evidence that cortical cells can respond with high temporal precision and reliability relative to a stimulus, and therefore relative to each other as well.

Deep-layer LEP spectra showed that cortical state spontaneously switched between two extremes: the synchronized and desynchronized state (Fig. 1c, Fig. 2a—e). There are many non-perceptual tasks that even primary sensory cortices might be engaged in during stimulus presentation, including attention (Roelfsema et al., 1998), memory formation and recall (Ji and Wilson, 2007), reward encoding (Shuler and Bear, 2006), locomotion (Saleem et al., 2013), visualization (Kosslyn et al., 1999), synaptic renormalization (Turrigiano et al., 1998), and cellular maintenance (Vyazovskiy and Harris, 2013). Many of these tasks have little to do with encoding the currently presented stimulus. To deal with this multitude of tasks, cortex may need to perform task switching, which could be reflected in cortical state changes.

Single unit responses to natural scene movie clips were more precise, reliable and sparse in the synchronized than desynchronized state (Fig. 6). The same held for LFP and MUA responses (Fig. 10), showing consistency across measures and types of signals. This result is surprising, because it conflicts with recent studies in V1 (Goard and Dan, 2009), primary auditory cortex (A1) (Marguet and Harris, 2011; Pachitariu et al., 2015) and primary somatosensory cortex (S1) (Hirata and Castro-Alamancos, 2011; Zagha et al., 2013) of anesthetized rodents. These studies come to the opposite conclusion: responses are more precise and reliable in the *desynchronized* state.

Several experimental differences might explain this conflicting result: differences in species (cat vs. rodent), anesthetic (isoflurane vs. urethane, ketamine/xylazine and fentanyl/medetomidine/-midazolam), desynchronization method (spontaneous vs. evoked), cortical area (V1 vs. A1 and S1), stimulus modality (visual vs. auditory and tactile), stimulus type (naturalistic vs. reduced), and the use of movie saccades. Since cortical state is likely multidimensional and SI measures only one such dimension (Harris and Thiele, 2011), it is also possible that there were other undetected changes in cortical state in the results presented here but not in those reported in the literature (or vice versa). Such undetected changes might account for some of these opposing results.

The species difference may be the most important. Cats have greater columnar organization of stimulus features in V1 than do rodents, including ocular dominance and orientation columns that rodents lack (Horton and Adams, 2005). UP phases in the synchronized state can manifest as waves of spontaneous activity travelling across the cortical surface (Petersen et al., 2003; Massimini et al., 2004; Benucci et al., 2007; Luczak et al., 2007; Xu et al., 2007; Mohajerani et al., 2010; Sato et al., 2012), while oriented visual stimuli can evoke standing waves of activity aligned to orientation columns (Benucci et al., 2007). Presumably, stimulus-evoked standing waves are absent in species that lack orientation columns, including rodents. Perhaps an interaction between travelling and standing waves of activity in the synchronized state increases the temporal precision and reliability of stimulus-evoked responses in cat but not rodent V1. This hypothesis predicts that responses in anesthetized ferret and primate V1, which also have orientation columns, should also be more precise and reliable in the synchronized state. Conversely, if there is a similar amount of stimulus feature map organization in A1 and S1 of both rodents and higher mammals (i.e., less than in V1 of higher mammals), this hypothesis also predicts that responses of anesthetized cat, ferret and primate A1 and S1 will be more precise and reliable in the desynchronized state, as is the case in rodents (Marguet and Harris, 2011; Hirata and Castro-Alamancos, 2011; Zagha et al., 2013; Pachitariu et al., 2015). This hypothesis may also provide an answer to the question of what functional role, if any, cortical columns might play (Horton and Adams, 2005): to increase response precision,reliability and sparseness. Further experiments that specifically take cortical state into account in sensory areas of anesthetized higher mammals in response to naturalistic stimulation are required to test these predictions.

More broadly, our results also conflict with the general understanding that responses in awake animals are enhanced during attending behavior (when cortex is more desynchronized) compared to quiescent resting behavior (when cortex is more synchronized) (Roelfsema et al., 1998; Fries et al., 2001; Cohen and Maunsell, 2009; Mitchell et al., 2009; Chalk et al., 2010; Pinto et al., 2013; Reimer et al., 2014). Our results therefore conflict with the hypothesis that synchronized and desynchronized cortical states in anesthetized animals are respectively analogous to quiescent and attending periods in awake animals (Luczak et al., 2007; Harris and Thiele, 2011; Luczak et al., 2013). Perhaps the relationship is more complex than previously thought. Indeed, some studies have suggested that the relationship between brain state, behavioral state, and the fidelity of stimulus representation can be surprisingly complex (Wikler, 1952; Podvoll and Goodman, 1967; Bradley, 1968; Sachidhanandam et al., 2013; Tan et al., 2014). Alternatively, periods of awake but unattending behavior may not be directly comparable to the globally synchronized state in anesthetized animals because the awake animal may still be attending to something else outside of the receptive fields of the recorded population. In other words, global vs. local synchronization (Vyazovskiy et al., 2011) under anesthesia vs. awake recordings, respectively, might help explain the inverted relationship between cortical state and response fidelity found here.

Although only indirectly shown here using global movie motion (Fig. 11), higher precision and reliability of responses during the synchronized state suggest that stimuli are better encoded, and hence more easily decoded, in the synchronized state. Why? With more numerous response events, narrower response events that are less likely to overlap with one another in time, and greater reliability of response events across trials, spike trains in the synchronized state are more distinctive than in the desynchronized state (Fig.3–Fig. 5), and should therefore be easier to decode. This has been shown more explicitly in other studies (Goard and Dan, 2009; Pachitariu et al., 2015), but with the opposite conclusion regarding cortical state.

The synchronized and desynchronized cortical states are two ends of a spectrum (Harris and Thiele, 2011; Luczak et al., 2013), and represent perhaps the simplest division of recording periods into different states. The synchronized state is itself composed of rapidly alternating up and DOWN phases, and the frequency content of the desynchronized state can be highly heterogeneous (Fig. 1c, Fig. 2a—e). A more thorough characterization of especially the desynchronized state is needed. Perhaps it may cluster into one of several sub-states (Gervasoni et al., 2004). More detailed characterization of brain state may reveal further surprises among neural responses.

## Acknowledgements

This work was supported by grants from the Canadian Institutes of Health Research and the Natural Sciences and Engineering Research Council of Canada. We thank Curtis L.Baker and Artur Luczak for detailed comments on the manuscript.

### Author Contributions

M.A.S. and N.V.S. conceived of and performed the experiments. M.A.S. analyzed the data and wrote the manuscript.

